# Aspects of invasiveness of *Ludwigia* and *Nelumbo* in shallow temperate fluvial lakes

**DOI:** 10.1101/504084

**Authors:** Viktor R. Tóth, Paolo Villa, Monica Pinardi, Mariano Bresciani

**Author notes:** **Correspondence:** Viktor R. Tóth Paolo Villa.

## Abstract

The relationship between invasive plant functional traits and their invasiveness is still the subject of scientific investigation, and the backgrounds of transition from non-native to invasive species in ecosystems are therefore poorly understood. Furthermore, our current knowledge on species invasiveness is heavily biased toward terrestrial species; we know much less about the influence of allochthonous plant traits on their invasiveness in aquatic ecosystems. We studied physiological and ecological traits of two introduced and three native macrophyte species in the Mantua lakes system (northern Italy). We compared their photophysiology, pigment content, leaf reflectance, and phenology in order to assess how the invasive *Nelumbo nucifera* and *Ludwigia hexapetala* perform compared to native species, *Nuphar lutea, Nymphaea alba*, and *Trapa natans*. We found *L. hexapetala* to have higher photosynthetic efficiency and able to tolerate higher light intensities than *N. nucifera* and the native species especially at extreme weather conditions (prolonged exposure to high light and higher temperatures). Chlorophyll a and b, and carotenoid contents of both allochthonous species was substantially higher than that of the native plants suggesting adaptive response to the ecosystem of Mantua lakes system. Higher variability of recorded data in invasive species also was observed. These observations suggest advanced photosynthetic efficiency of the invasive species, especially *L. hexapetala*, resulting in faster growth rates and higher productivity. This was supported by the evaluation of seasonal dynamics mapped from satellite remote sensing data. This study provides empirical evidence for the relationship between specific plant physiological traits and invasiveness of aquatic plant species, highlighting the importance of trait studies in predicting ecosystem-level impacts of invasive plant species.

## 1 Introduction

Introduced species pose an imminent threat to biodiversity, species composition and structure, as well as general functioning of their new ecosystems (D’Antonio and Kark, 2002; Vilà et al., 2011; Vitousek, 1990). This is especially the case in aquatic environments and for aquatic plants, or macrophytes (Gallardo et al., 2016). Invasive species are responsible for displacing large numbers of native species throughout the world (Holmes and Cowling, 1997; Meyer and Florence, 1996; Musil, 1993). Moreover, the consequences of invasions are long-term; otherwise inaccessible regions of the planet maintain new invasive populations that are practically inerasable (Coblentz, 1990). Although the introduction of species to new environments can result from natural processes, the vast majority in recent decades has occurred as either a direct (i.e., purposeful transport) or indirect (i.e., without human awareness) consequence of the anthropogenic collapse of biogeographic barriers (McKinney and Lockwood, 1999). In addition, anthropogenic pressure on ecosystems (e.g., altered land-use patterns, changes in climate regimes, increase in atmospheric CO_2_), and especially on aquatic ecosystems, foster favourable conditions for the establishment of introduced species (Dukes and Mooney, 1999; Fasoli et al., 2018; Hussner et al., 2014).

Multiple differences between native and introduced species have been observed, and can be summarized as being associated with either more efficient resource use by the invasive plants (Baruch and Goldstein, 1999; Vitousek, 1986), or with invasive species’ lack of natural enemies in their new habitats (Callaway and Aschehoug, 2000; Mitchell and Power, 2003). Despite the advantages the introduced species have, this does not always correspond to biomass dominance; indeed, the background processes of how these advantages are translated into better performance in their new environments are still not fully understood (Davis et al., 2001; Funk and Vitousek, 2007). Even though invasive plants seem to have higher plasticity and enhanced functional traits in many instances (Davidson et al., 2011; Pyšek and Richardson, 2007), the recent scientific literature has shown no consensus as to whether the carbon acquisition and distribution of native and invasive plants differ (Daehler, 2003; Leffler et al., 2014; Van Kleunen et al., 2010).

Within this context, macrophytes emerge as particularly interesting targets, for several reasons. Macrophytes tend to display higher diversity in temperate areas (Alahuhta et al., 2017), where human impacts are stronger, to show highly cosmopolitan features (Zhou and Zhou, 2009), and to occupy the extremes of the global spectrum of vegetation forms (Díaz et al., 2016). As such, macrophytes are more sensitive to ecosystem degradation and invasion processes, but also provide insight into the functional adaptation potential of plants to environmental conditions different from those of their native range.

The main aim of this study was to investigate and compare selected traits (physiological, spectral, and seasonal dynamics) of invasive and native macrophyte species found in a shallow temperate freshwater system. Towards this, we hypothesised that: i) photophysiological performance (i.e., chlorophyll fluorescence, related to C acquisition) of invasive species is superior to that of native plants; ii) leaf pigment pools of invasive plants are different in size and composition from native ones; iii) photophysiological performance and pigment content show higher plasticity in invasive than in native species; iv) invasive macrophytes can effectively exploit temporal niches left unoccupied by the seasonal dynamics of native species; and that v) leaf spectroscopy can provide information about physiological traits of both invasive and native macrophytes.

## 2 Materials and Methods

### 2.1 Study sites

The Mantua lakes system (Laghi di Mantova), consisting of the Superior, Middle, and Inferior lakes, are three shallow (3.5 m mean depth) fluvial lakes adjacent to the city of Mantua, Italy (Fig. 1). The lakes are surrounded by urban areas, both residential and industrial, and the upstream basin is characterized by intense agricultural cultivation. The Mantua lakes system is therefore exposed to diffuse sources of nutrients (N and P) and other pollutants. As a result, the lakes system is characterized by high turbidity (Secchi disk depth < 1 m in summer), high trophic level (Chl-a up to 200 mg m^-3^), and the coexistence of phytoplankton and predominantly the following macrophytes, comprising emergent, submerged, floating-leaved and free-floating species (Bolpagni et al., 2014; Bresciani et al., 2013; Pinardi et al., 2011; Villa et al., 2015).

**Figure 1.**
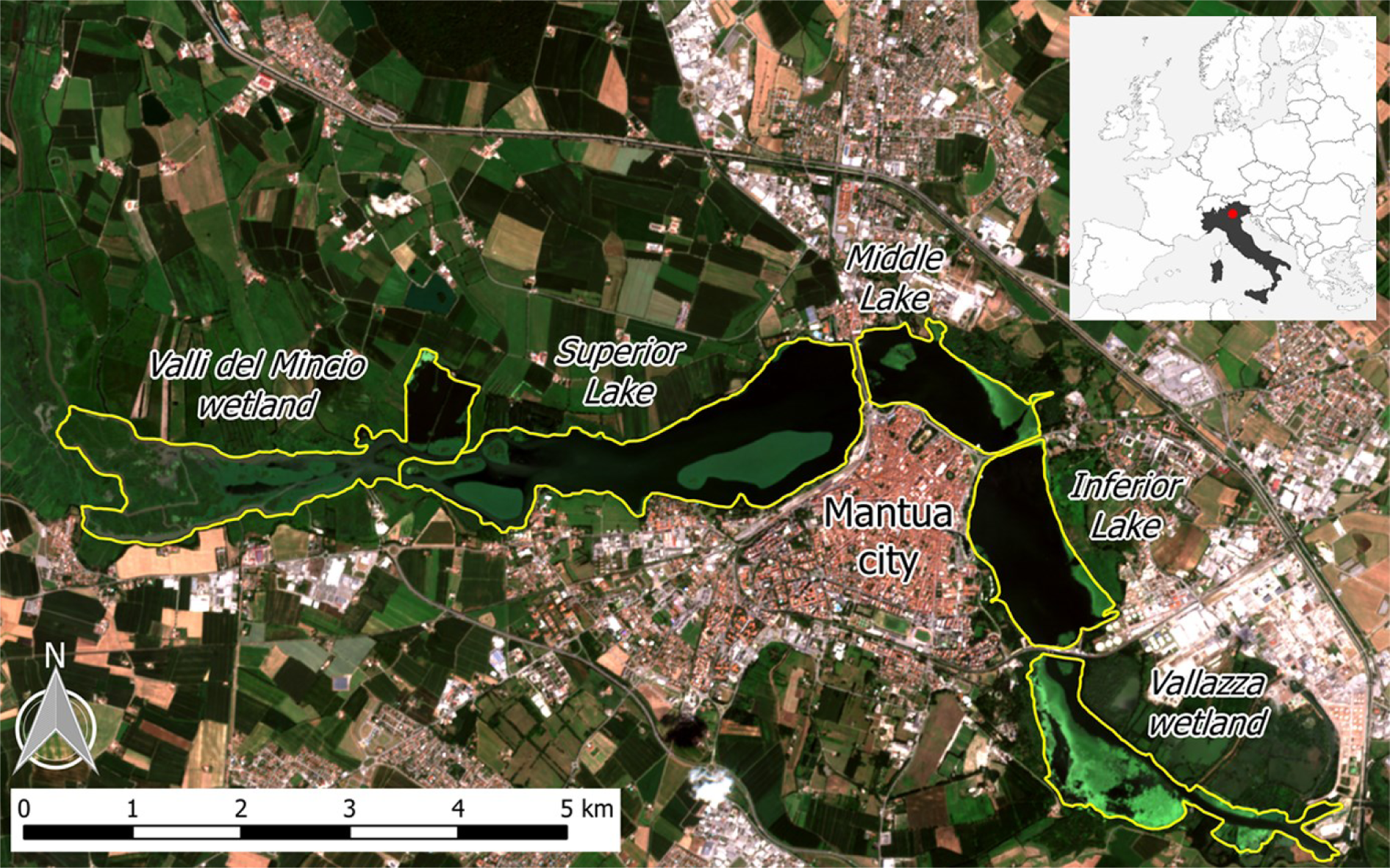
The Mantua lakes system, with the location of the study area in Italy (upper right box), and the different subsystems (lakes and wetlands) indicated. Background image is a Sentinel-2A satellite-acquired true colour composite from 28 July 2016.

The sacred lotus (*Nelumbo nucifera* Gaertn.), an allochthonous species originating from Southeast Asia, colonizes Superior Lake, forming two large macrophyte islands. The plant is rooted in the sediment and can grow in waters up to 3 m deep. The large, shield-shaped leaves ascend from the rhizomes on waxy, prickly stems and can rise above the water level. Various parts of the plant can be used for different purposes, including food, silk production, and medicine, as well as for esthetical reasons, mainly due to its large, beautiful flowers. *Nelumbo* was originally introduced to the Mantua lakes system in the 1920s, with its rhizomes proposed as an alternative food source. In the subsequent decades, it has been mainly been used for floral collection, and has spread throughout the lakes over the past five decades.

The water primrose (*Ludwigia hexapetala* (Hook. & Arn.) Zardini, H.Y. Gu & P.H. Raven), native to South America, is the other allochthonous species that has spread in the lakes over the past decade, first colonizing calm water bays in the Middle Lake and more recently the riparian zones of Superior and Inferior lakes. In its native range, *Ludwigia* can be found in wet grasslands and wetlands, whereas in areas where it has been introduced, it is able to invade lakes, slow fluvial plumes, and wetlands, and is one of the most invasive plant genera in Europe (EPPO, 2004b, 2004a). *Ludwigia* has large (2-5 cm), yellow flowers, and 8-10 cm long, spear-shaped, pilose leaves. The stem of the plant grows from 2-3 meter-deep water to the water surface, and can emerge above the water level.

Water chestnut (*Trapa natans* L.) is the most widespread native species, present in all three lakes and dominant in Middle and Inferior lakes. Nymphaeids, mainly *Nuphar Lutea* (L.) Sm *Nymphaea alba* L., are present in the upstream portion of Superior Lake, as well as in small patches in Middle and Inferior lakes. Submersed species (e.g., *Ceratophyllum demersum* L., *Vallisneria spiralis* L., and *Najas marina* L.), predominantly *C. demersum*, are especially present in Superior Lake.

The lakes system is part of the protected Mincio Regional Park, which manages water and macrophytes. The system is characterized by high nutrient and organic matter loads and low water flow rates, resulting in its tendency towards infilling and heightened risk of hypoxia (Bolpagni et al., 2014; Pinardi et al., 2011). For this reason, since 2004, *N. nucifera* and *T. natans* macrophyte stands are controlled by cutting and occasionally harvesting (Pinardi et al., 2011; Villa et al., 2017).

### 2.2 Photophysiological measurements

Chlorophyll fluorescence parameters were measured using a chlorophyll fluorometer (PAM-2500, Heinz Walz GmbH, Germany) between 9h00 and 15h00. Light response curves (i.e., the electron transport rate (ETR) of the photosystem II (PSII), as a function of photosynthetically active radiation (PAR)) were measured from mature, healthy-looking leaves after a dark-adapting period of 20 minutes. After dark adaptation, emitted initial fluorescence yield (F_o_) and maximal fluorescence yield (F_m_) resulting from a pulse of a saturated light (630 nm, intensity 3000 μmol m^−2^ s^−1^) were determined. From these, the photochemical PSII efficiency (F_v_/F_m_), coefficient of photochemical quenching (qP), and coefficient of non-photochemical quenching (qN) were calculated. The measured leaves were exposed to 11 actinic lights for a duration of 15 seconds, at 630 nm, with an intensity of between 5 and 787 μmol m^−2^ s^−1^, and the ETR values were measured after each illumination step with a new pulse of saturated (3000 µmol m^−2^ s^−1^) light. Exponentially saturating curves (Eilers and Peeters, 1988) were fit to the light response data, and the maximum ETR (ETR_max_), theoretical saturation light intensity (I_k_), and maximum quantum yield for whole chain electron transport (α) were retrieved using this formula (Genty et al., 1989).

### 2.3 Pigment analysis

Leaf discs of 0.6 cm in diameter, were collected from intact leaves close to where chlorophyll fluorescence was measured, and stored in aluminium foil in sub 0 temperature until their transfer to a –20 °C freezer. Frozen leaf discs samples were homogenised in liquid N_2_, then extracted in 80% acetone. The extracts were centrifuged and the supernatant collected and stored at –20 °C until analyses. The absorbance spectra (400-750 nm) of extracts were measured using a spectrophotometer (Shimadzu UV-2401PC, dual-beam), and pigment concentrations (chlorophyll-a (Chl-a), chlorophyll-b (Chl-b), and total carotenoids (Car) were calculated using empirical formulae (Wellburn, 1994) and reported as fresh weight concentrations (µg g_fw_^-1^).

### 2.4 Spectroradiometric measurements

Leaf reflectance of the same leaves sampled for photophysiological measurements was measured over the visible to shortwave infrared spectral range (350 – 2500 nm) using a portable SR-3500 Full Range spectroradiometer (Spectral Evolution, Lawrence, USA). Following a 20-minute dark adaptation period, leaf-reflected radiance under near-steady state conditions (60 seconds after leaf de-shadowing) was measured using a Reflectance Contact Probe (Spectral Evolution, Lawrence, USA). The probe was equipped with a light source (5-watt tungsten halogen bulb) and spectra were calibrated to leaf reflectance using measurements of a 99% Spectralon panel (Labsphere, North Sutton, USA). During measurements, macrophyte leaves were placed over a flat neoprene plate (absorbance > 95%) in order to minimize background reflection of light transmitted through the leaves. This approach was chosen because it is easier and faster under challenging field conditions (e.g., boat-based surveys) compared with using an integrating sphere to fully determine reflectance and transmittance, and only introduces minimal distortion in leaf reflectance measurements (Potůčková et al., 2016; Sims and Gamon, 2002).

Leaf reflectances were used to calculate a set of spectral indices (SIs), belonging to three different groups: (1) SIs sensitive to photosynthetic pigments (Gitelson et al., 2001, 2003; Sims and Gamon, 2002); (2) SIs connected to radiation use efficiency (RUE) and the state of the xanthophyll cycle pigments (Gamon et al., 1992; Garrity et al., 2011; Hernández-Clemente et al., 2011; Wu et al., 2010); and (3) SIs correlated to macrophyte, *Phragmites australis* (common reed), physiological parameters (Stratoulias et al., 2015). The SIs tested in this study, with their formulas and relevant references are provided in Supplementary Table 1.

**Table 1.** Coefficient of variation of the standardised pigment and photophysiological data of Mantua lakes system macrophytes. chl-a – chlorophyll a content; chl-b - chlorophyll b content; car – total carotenoids content; a/b - chlorophyll a to chlorophyll b ratio; chl/car - chlorophylls to carotenoids ratio; α - maximum quantum yield for whole chain electron transport; ETR_max_ - maximal electron transport rate; I_k_ - theoretical saturation light intensity; F_v_/F_m_ - PSII photochemical efficiency; qN - coefficient of non-photochemical quenching; and qP - coefficient of photochemical quenching.

| Species | chl-a | chl-b | car | a/b | chl/car | $\alpha$ | $ETR_{max}$ | $I_k$ | $F_v/F_m$ | qN | qP |
| --- | --- | --- | --- | --- | --- | --- | --- | --- | --- | --- | --- |
| *Ludwigia* | 0.26 | 0.46 | 0.18 | 0.84 | 0.35 | 1.14 | 3.11 | 1.41 | 0.75 | 0.83 | 0.36 |
| *Nelumbo* | 0.41 | 0.40 | 0.33 | 0.31 | 0.24 | 0.34 | 0.04 | 0.15 | 0.53 | 0.70 | 0.35 |
| *Nuphar* | 0.05 | 0.09 | 0.08 | 1.08 | 0.22 | 1.66 | 0.05 | 0.20 | 1.03 | 0.57 | 1.10 |
| *Nymphaea* | 0.02 | 0.03 | 0.05 | 0.41 | 0.08 | 0.38 | 0.01 | 0.12 | 0.78 | 0.40 | 0.37 |
| *Trapa* | 0.04 | 0.07 | 0.04 | 0.75 | 0.56 | 0.93 | 0.05 | 0.21 | 1.35 | 1.11 | 1.28 |

In order to focus on the most relevant leaf spectral features, the collinearity of all SIs tested was assessed by computing the correlation coefficient (Pearson’s *r*) of each SI pair (Supplementary Fig. 1). Only those SIs carrying the most information were retained for further analysis and discussion, by excluding the ones those with *r* > 0.9.

### 2.5 Summary of *in situ* measurements

Leaf samples were measured in the Mantua lakes system in late July, 2016, and late May and late July, 2017. A summary of sample sites and number of samples is provided in Supplementary Table 2. Leaves were collected from plants located within 2-3 m of the water edge of homogenous, intact macrophyte stands. For *in situ* measurements, from each plant the youngest, mature, intact leaf was chosen. At each sampling site, 3 to 12 plants per species were selected (Supplementary Table 2).

Following dark adaptation, whole leaves (for *Ludwigia* and *Trapa*) or leaf parts (for *Nelumbo, Nuphar* and *Nymphaea*) were removed from the plants and subjected to measurements. Chlorophyll fluorescence measurements were performed on the dark-adapted part of the leaves, with leaf spectroradiometric reflectance recorded in its close vicinity. In order to avoid possible bias due to light environment differences, only leaves exposed directly to sunlight were measured.

### 2.6 Seasonal dynamics

Additional information highlighting key macrophyte phenological characteristics were extracted from macrophyte LAI seasonal dynamics maps. Time series of LAI for floating and emergent macrophytes in the Mantua lakes system were derived from medium resolution satellite data (Landsat 8, SPOT5, and Sentinel-2) for the 2015 growing season (Villa et al., 2018). From these time series, seasonal dynamics metrics were computed, namely: the day of the start of season (SoS), the day of the peak of season (PoS), the day of the end of season (EoS), the duration of the growing season (Length), the maximum LAI value (LAI_max_), the rate of increase of LAI during the early growth (Growth rate), and the rate of decrease of LAI during the senescence (Senescence rate).

We calculated the mean and standard deviation scores of these seasonality metrics of all pixels dominated by each macrophyte species considered in this work, as well as the area covered by each species and the corresponding percentage of total macrophyte cover. *Nymphaea* pixels are grouped together with *Nuphar* pixels, because they only cover a very small area (<1 ha).

### 2.7 Statistical analyses

Graphing, curve fitting, and statistics were performed in Past (Hammer et al., 2001) and various packages in R v.3.4.4 (R Development Core Team, 2012).

## 3 Results

### 3.1 Photophysiological measurements

Clear differences in the photophysiological properties of the studied plants and in the photosynthetic activity of *Ludwigia* were observed (Supplementary Figure 2; Fig. 2). Not only did the extent of ETR_max_ and I_k_ differ significantly for *Ludwigia* with respect to the rest of the species (Kruskal-Wallis One Way Analysis of Variance on Ranks; H=57.2; P<0.001; n=182), but their distributions were bimodal, suggesting the presence of two subpopulations (Fig. 2). These separate *Ludwigia* subpopulations can partially be explained by the marked differences between 2016 and 2017 samples: ETR_max_ scores for *Ludwigia* are in fact significantly lower in July 2016 than in July 2017, 162.2±15.9 and 743.4±290.2 µmol m^-2^ s^-1^ respectively (Mann–Whitney *U* test; *U*=0; P<0.001; n_1_=n_2_=9). Due to this significant difference in ETR_max_ and in I_k_, the position of *Ludwigia* in the photophysiological PCA was well defined (Fig. 3).

**Figure 2.**
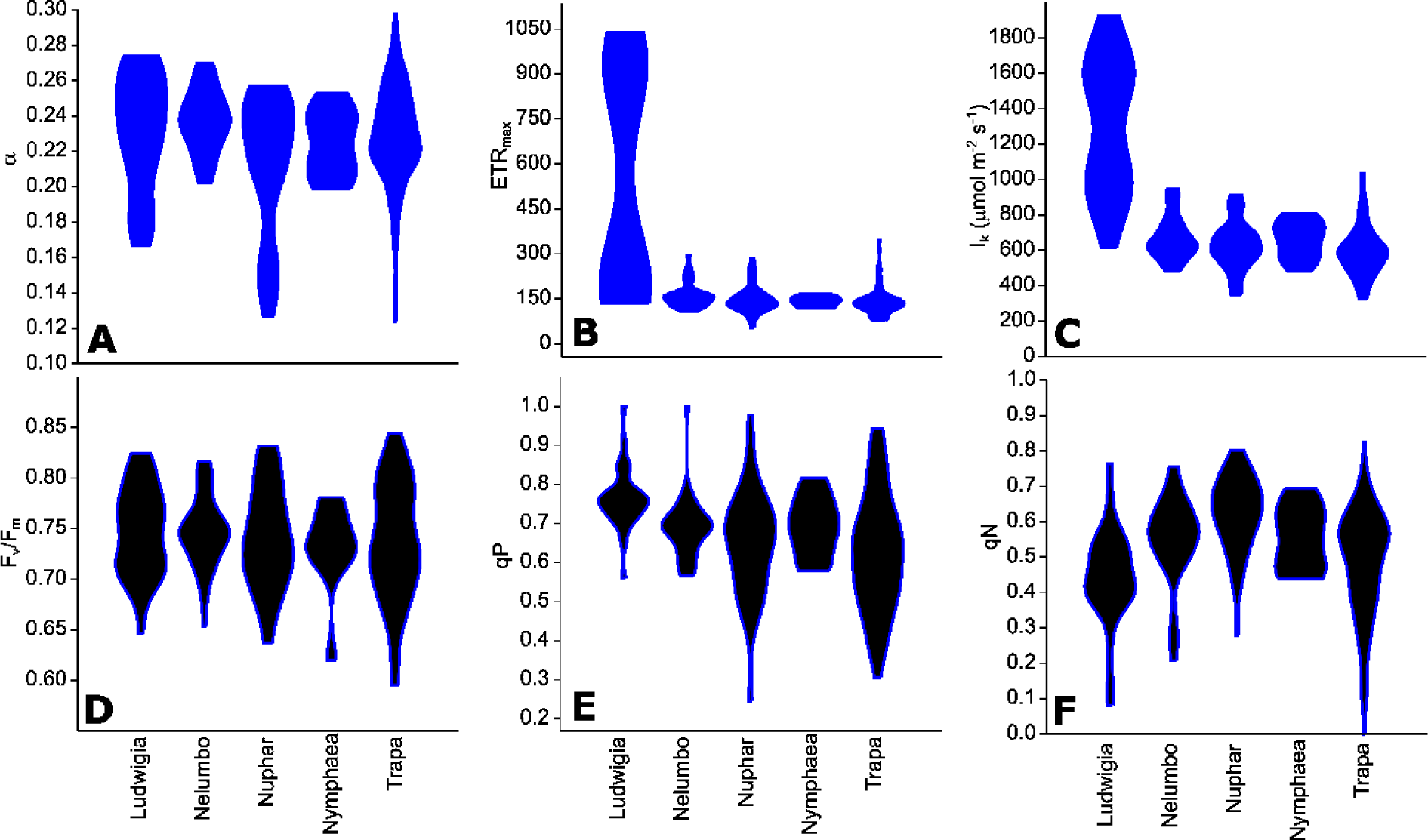
Chlorophyll fluorescence parameters measured from Mantua lakes system macrophytes. A - maximum quantum yield for whole chain electron transport (α), B - maximal electron transport rate (ETR_max_), C - theoretical saturation light intensity (I_k_), D - PSII photochemical efficiency (F_v_/F_m_), E - coefficient of photochemical quenching (qP), and F - coefficient of non-photochemical quenching (qN).

**Figure 3.**
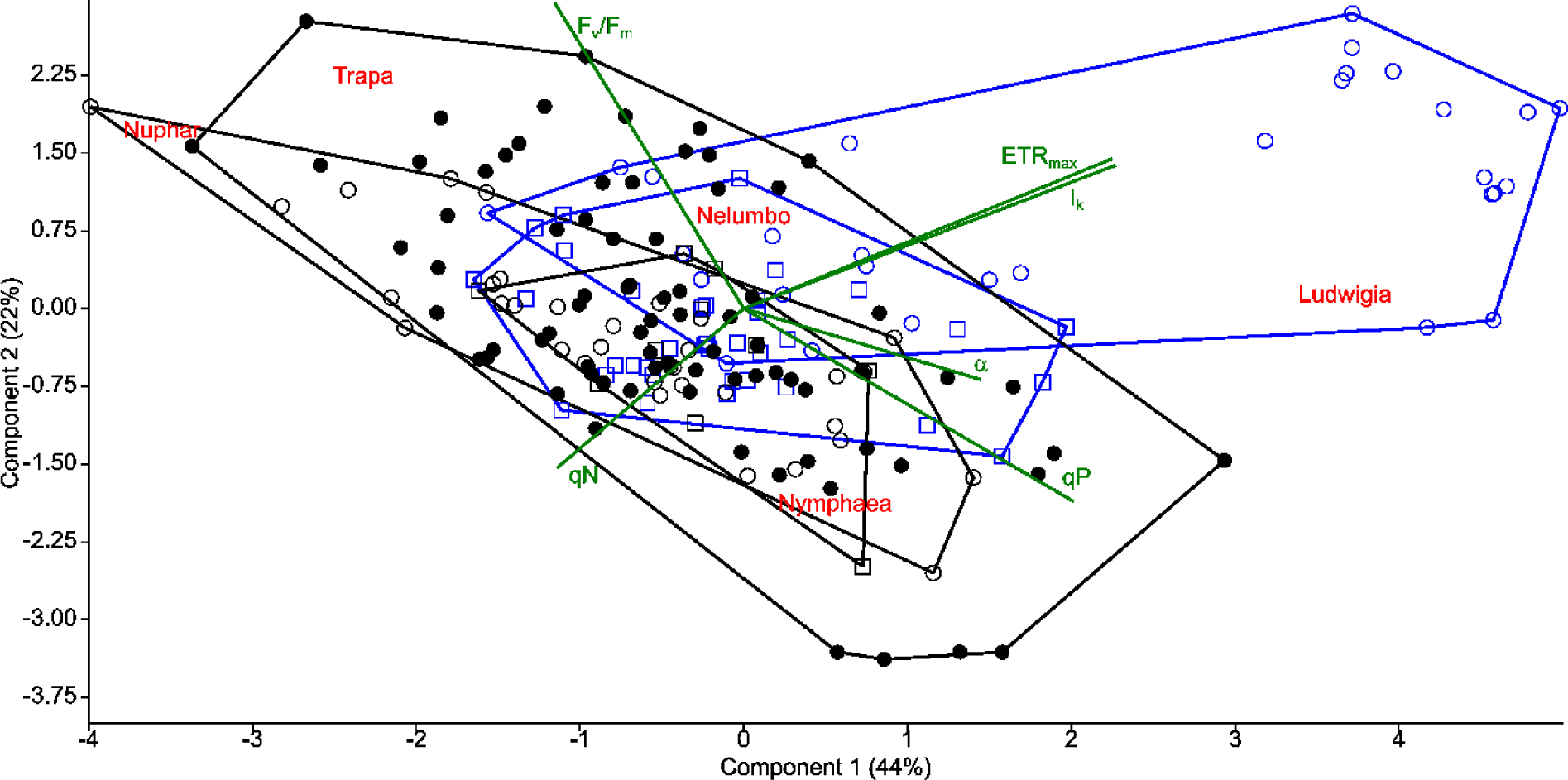
Principal component analysis (PCA) of photophysiological parameters measured from macrophytes of the Mantua lakes system. α - maximum quantum yield for whole chain electron transport, ETR_max_ - maximal electron transport rate, I_k_ - theoretical saturation light intensity, F_v_/F_m_ - PSII photochemical efficiency, qP - coefficient of photochemical quenching, and qN - coefficient of non-photochemical quenching.

The photophysiological properties of *Nelumbo* did not differ from those of the local macrophytes, therefore *Nelumbo* aligned with the rest of the species present within the PCA. Further analysis of the photophysiological parameters of the studied macrophytes demonstrated the higher plasticity of *Ludwigia, Trapa*, and *Nuphar* compared with *Nymphaea* and *Nelumbo* (Table. 1).

### 3.2 Pigment analysis

Chl-a, Chl-b, and Car content of *Nelumbo* and *Ludwigia* were significantly higher than those of other species (Figs. 4A, 4B and 4C; Kruskal-Wallis One Way Analysis of Variance on Ranks; P<0.001). *Ludwigia* Chl-a content was higher in May than in July, 2001±258 compared with 1591±313 µg g_fw_ ^-1^ (Mann–Whitney *U* test; *U*=11; P=0.01; n_1_=n_2_=9). Nevertheless, despite the higher pigment content - *Ludwigia* and *Nelumbo* had, on average, 2.8, 2.7, and 2.4 times higher Chl-a, Chl-b, and Car content than *Nymphaea, Nuphar*, and *Trapa* respectively - the Chl-a/ Chl-b ratio (a/b) of *Ludwigia* alone differed significantly (P < 0.05) from that of *Nelumbo, Nuphar*, and *Nymphaea*, and was similar to that of *Trapa*. The total chlorophyll (Chl-a + Chl-b) to Car ratio (chl/car) of *Nymphaea* was significantly lower (2.18±0.05) than the rest of the species (3.56±0.05).

**Figure 4.**
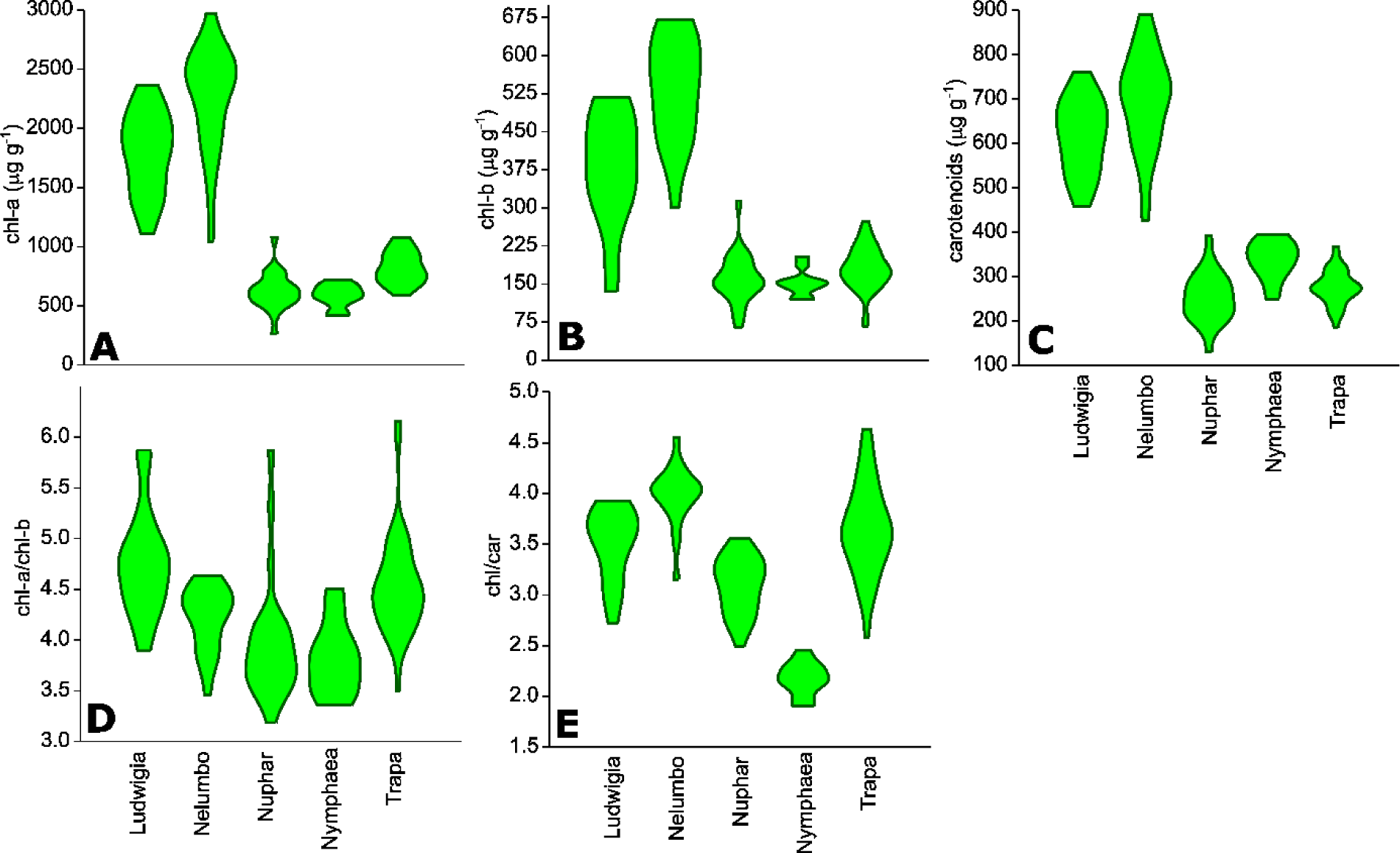
Leaf pigment contents of macrophytes of the Mantua lakes system, reported as a function of fresh weight. A –chlorophyll a content (chl-a; μg g_fw_^-1^), B - chlorophyll b content (chl-b; μg g_fw_^-1^), C – total carotenoids content (carotenoids; μg g_fw_^-1^), D – chlorophyll a to chlorophyll b ratio (chl-a/chl-b), E – chlorophylls to carotenoids ratio (chl/car).

Due to the significant difference of *Ludwigia* and *Nelumbo* from native species in leaf pigments content, their discrimination was clearly defined through PCA (Fig. 5). Further analysis of pigment content results showed high phenotypic plasticity of Chl-a, Chl-b, and Car content for both *Ludwigia* and *Nelumbo*, although pigment stoichiometry was also variable for *Nuphar* and *Trapa* (Table. 1).

**Figure 5.**
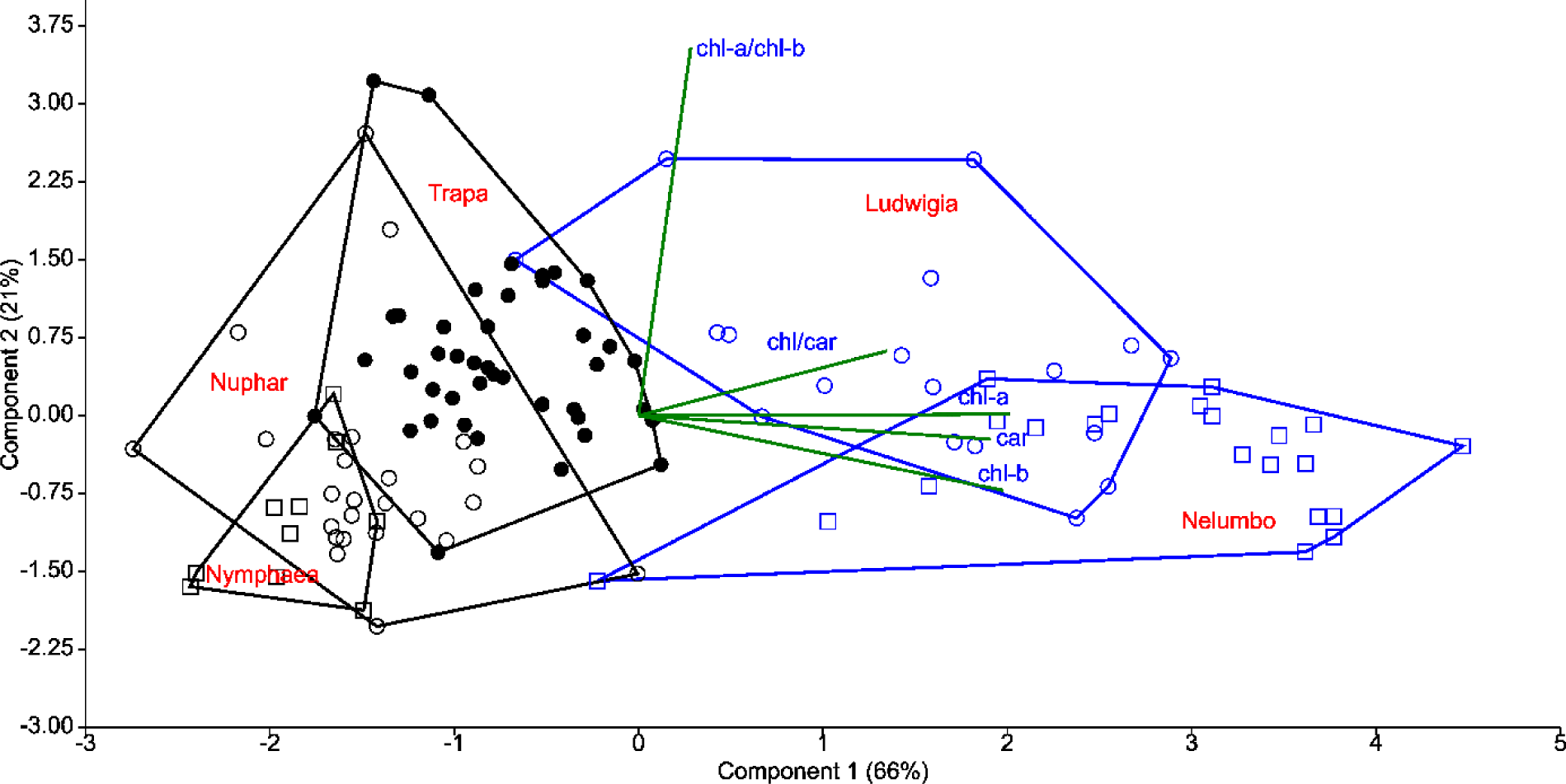
Principal component analysis (PCA) of leaf pigment content of macrophytes of the Mantua lakes system. chl-a – chlorophyll a content; chl-b - chlorophyll b content; car – total carotenoids content; chl-a/chl-b - chlorophyll a to chlorophyll b ratio; and chl/car - chlorophylls to carotenoids ratio.

### 3.3 Spectroradiometric measurements

After assessing the cross-correlation of tested SIs of the sampled species (Supplementary Fig. 1), some indices were excluded from further analyses. The list of SIs tested, with their full names and relevant references are included in Supplementary Table 1. PRI_r2_ and PRI*CI were excluded because of their high correlation with PRI (*r* = 0.94 and *r* = 0.95 respectively). ND_539,560_ was also not considered further due to its high correlation with PRI (*r* = 0.92). Within the macrophyte eco-physiological parameter SIs, ND_659,687_ and ND_621,692_ were found to be highly correlated (*r* = 0.91), so we therefore only retained the latter for further analysis. Finally, we excluded ND_705_ due to its high correlation with mND_705_ (*r* = 0.84), and CI due to its lower sensitivity to leaf chlorophyll content compared to mND_705_ (*r* = 0.41-0.42 and *r* = 0.77-0.78 respectively).

Within our dataset, there is a significant difference in PRI between the species (Kruskal-Wallis One Way Analysis of Variance on Ranks; H=45.8; P <0.001; n=152). PRI of *Nelumbo* samples was higher (Dunn’s Multiple Comparison test, P<0.05) than that of *Trapa, Nuphar*, and *Nymphaea*, while *Ludwigia* was not found to be significantly different, although on average PRI was lower. PRI_515_ differs from PRI especially for the allochthonous species, and in particular for *Nelumbo*, which shows significantly lower PRI_515_ scores compared to the autochthonous species (Dunn’s Multiple Comparison test, P<0.05).

Our PRI observations generally concur with previous findings of which derived indices are best spectral proxies for aquatic plant physiological parameters (Stratoulias et al., 2015). *Nelumbo* is statistically different from all the other species (Dunn’s Multiple Comparison test; P<0.05) in terms of ND_546,551_, and is different from the autochthonous species (Dunn’s Multiple Comparison test; P<0.05) in terms of ND_621,692_. Even though not statistically significant, ND_621,692_ scores for *Ludwigia* are also slightly lower than those of the autochthonous species (0.055±0.021 and 0.076±0.034 respectively, in July 2017).

**Figure 6.**
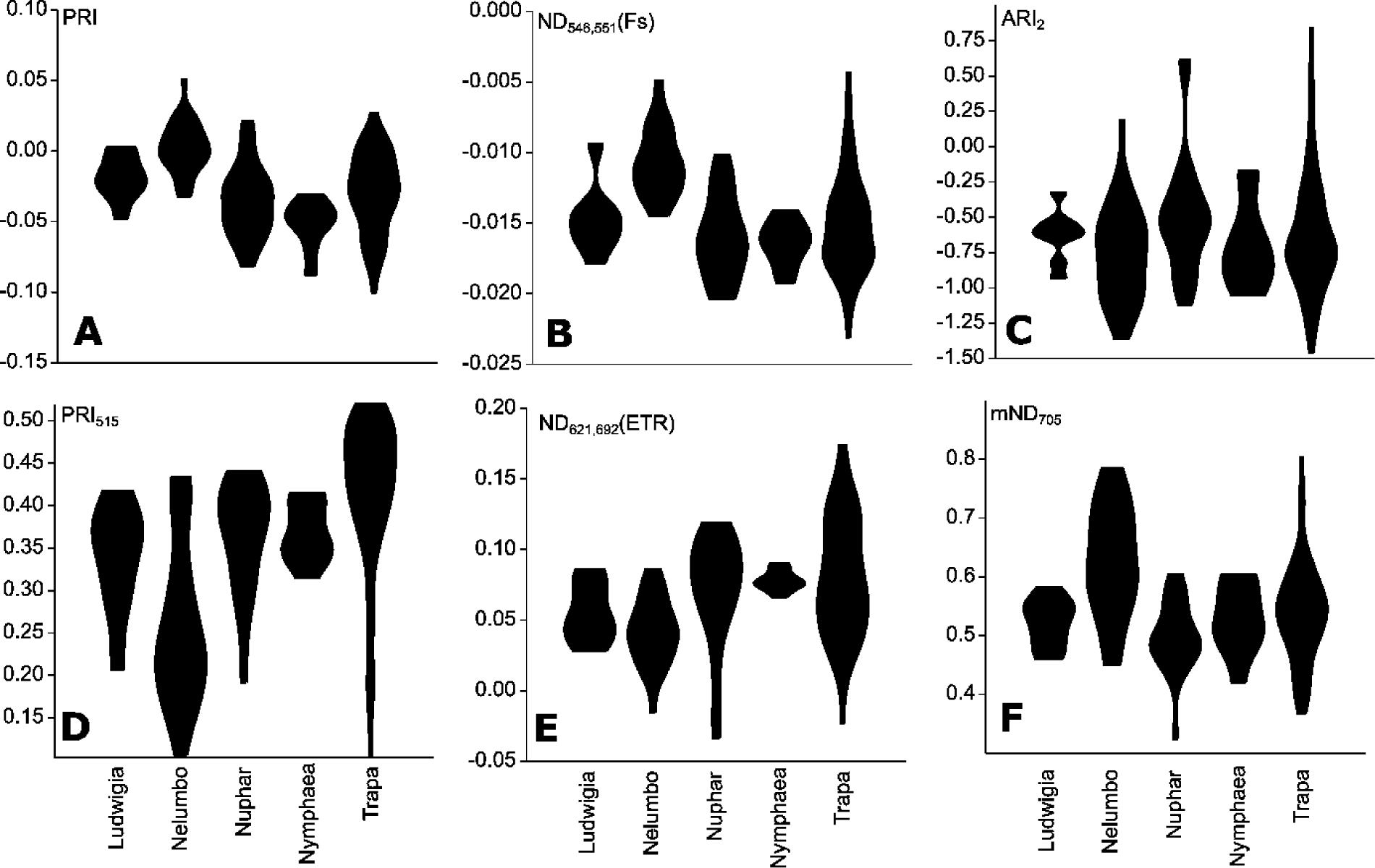
Selected SIs calculated from Mantua lakes system macrophyte leaf reflectance data of. A - Photochemical Reflectance Index (PRI); B - Normalized Difference Spectral Index 546,551 (ND_546,551_); C - Anthocyanin Reflectance Index 2 (ARI_2_); D - Photochemical Reflectance Index 515 (PRI_515_); E - Normalized Difference Spectral Index 621,692 (ND_621,692_); and F - modified Normalized Difference 750/705 (mND_705_).

**Figure 7.**
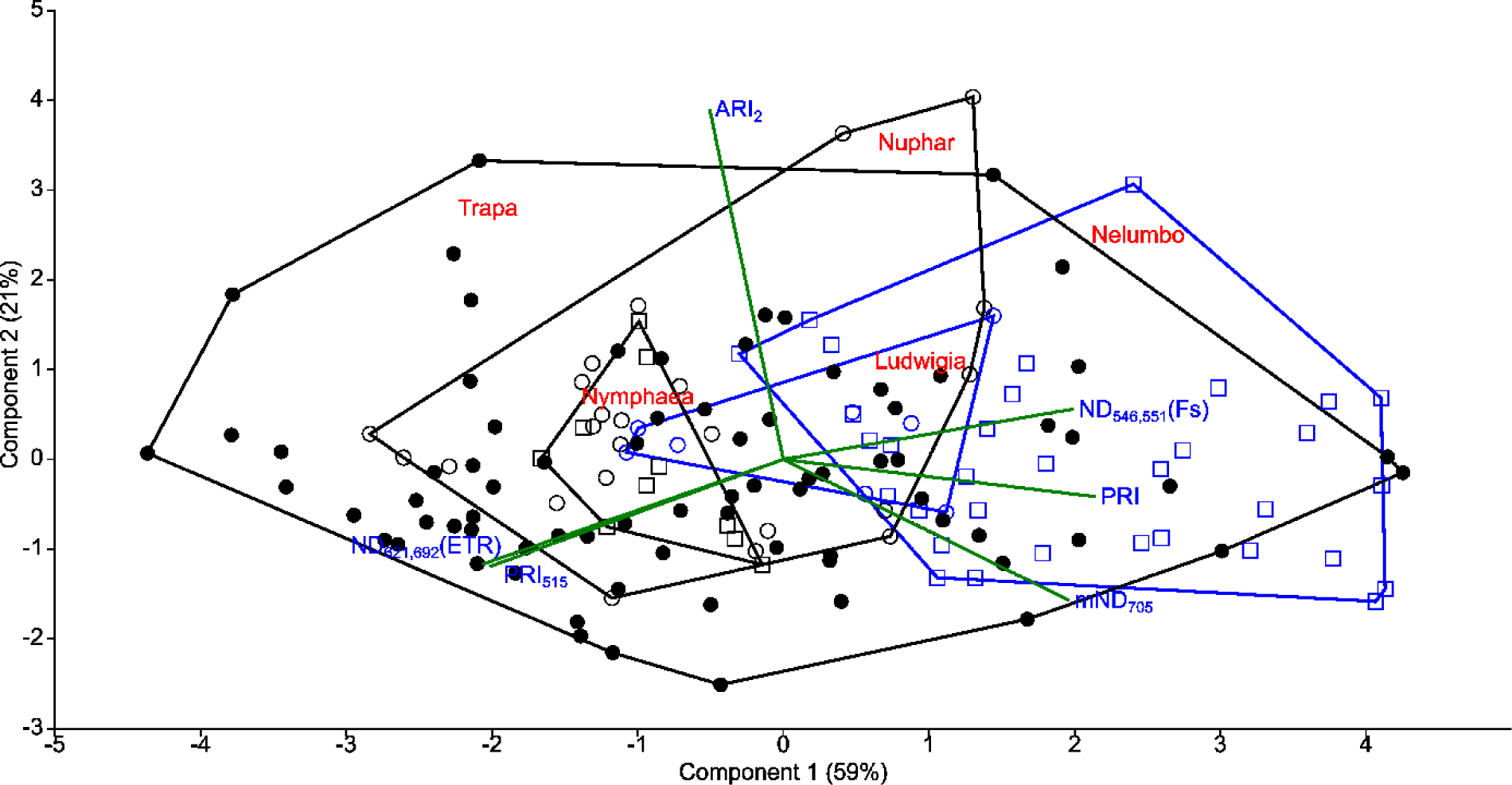
Principal component analysis (PCA) of selected SIs calculated from Mantua lakes system macrophyte leaf reflectance data. PRI - Photochemical Reflectance Index; ND_546,551_ - Normalized Difference Spectral Index 546,551; ARI_2_ - Anthocyanin Reflectance Index 2; PRI_515_ - Photochemical Reflectance Index 515; Normalized ND_621,692_ - Difference Spectral Index 621,692; and mND_705_ - modified Normalized Difference 750/705.

ARI_2_, which is related to secondary pigment contents, showed no difference between the species (Kruskal-Wallis One Way Analysis of Variance on Ranks; H=6.748; P=0.150; n=152). Rather, significant differences between the species are observed for mND_705_, related to total chlorophyll content (One Way Analysis of Variance; F=10.906; P<0.001; n=152), with *Nelumbo* showing higher values compared to any other species (post hoc Tukey test; P<0.008).

### 3.4 Seasonal dynamics features

Differences between allochthonous and autochthonous species, in terms of their seasonal dynamics, or phenology metrics (day of the start (SoS), peak (PoS), and end of season (EoS), as well as the length of growing season), are highlighted in Table 2, referring to 2015 growing season. Peak LAI values (LAI_max_) and LAI growth and senescence rates (LAI_growth_, LAI_senescence_) were calculated in our previous study (Villa et al., 2018), and are also reported.

**Table 2.** Synoptic metrics of seasonal dynamics (phenology and LAI growth) for Mantua lakes system macrophytes for 2015, derived from Villa et al. (2018). SoS – day of the start of season; PoS – day of the peak of season; EoS – day of the end of season; length - duration of the growing season (EoS-SoS); LAI_max_ - maximum LAI value; growth rate - rate of increase of LAI during the early growth; and senescence rate - rate of decrease of LAI during the senescence.

|  | *Ludwigia* | *Nelumbo* | *Nuphar-*<br>*Nymphaea* | *Trapa* |
| --- | --- | --- | --- | --- |
| SoS (DOY - Julian day) | 122.5±6.4 | 132.4±3.9 | 115.1±8.0 | 155.2±16.6 |
| PoS (DOY - Julian day) | 222.3±6.9 | 218.6±4.0 | 200.6±13.3 | 215.8±13.0 |
| EoS (DOY - Julian day) | 314.8±13.2 | 304.4±3.9 | 263.1±20.4 | 256.7±13.2 |
| Length (days) | 192.3±14.8 | 172.0±5.2 | 147.9±28.1 | 101.5±9.7 |
| LAI<sub>max</sub> (m<sup>2</sup> m<sup>-2</sup>) | 1.15±0.28 | 1.69±0.05 | 1.06±0.18 | 1.28±0.13 |
| Growth rate (m<sup>2</sup> m<sup>-2</sup> d<sup>-1</sup>) | 0.018±0.007 | 0.047±0.008 | 0.015±0.005 | 0.023±0.004 |
| Senescence rate (m<sup>2</sup> m<sup>-2</sup> d<sup>-1</sup>) | 0.013±0.004 | 0.026±0.004 | 0.018±0.006 | 0.045±0.018 |
| Area (ha) | 2 | 69 | 9 | 74 |
| % cover of total macrophytes | 1.3 | 44.8 | 5.8 | 48.1 |

The growing season of allochthonous species was found to be longer (192±15 days for *Ludwigia*, 172±5 days for *Nelumbo*) and to end later in the year (late October to mid-November) than for native species. *Nelumbo* also shows morphological advantages, with maximal leaf area index (LAI_max_) reaching 1.7 m^2^ m^-2^ at PoS, compared to 1.0-1.2 m^2^ m^-2^ for the other species, as well as fast early-season growth (i.e., LAI_growth_ – the velocity of LAI growth from SoS to PoS – is 2-3 times higher than that of the other species).

## 4 Discussion

The background and dynamics of invasion processes are still a matter of debate. Nevertheless, studies focusing on morphological and physiological characteristics of native and invasive plants can elucidate the success of the latter (Baruch and Goldstein, 1999; Hufbauer and Torchin, 2008; Van Kleunen et al., 2010). While a given explanation of how morphological and physiological parameters can promote the spread of non-native species might only be applicable to the ecosystem in question, some of the observations could be considered to be more generally valid. For example, higher photosynthetic rates of invasive plants could result in higher biomass accumulation, consequently exhausting resources available for native species and/or shading out native competitors (Baruch and Goldstein, 1999; Hufbauer and Torchin, 2008). Of all the species investigated, *Ludwigia* had the highest photosynthetic parameters (ETR_max_, I_k_), which is in line with its C_4_ nature and high photosynthesis rates (Dandelot et al., 2005; Hussner, 2009), although this does not always translate into higher biomass (Hussner, 2009). It is worth noting that in the Mantua lakes system, *Ludwigia* samples showed a strong response to environmental changes. The outstanding bimodality of ETR_max_ and I_k_ for this species could be explained by inter-seasonal differences between 2016 and 2017, linked to differences in meteorological conditions up to late July for the two years. When compared to 2016, 2017 was much drier (total precipitation 52% lower), hotter (average max temperature higher by 0.9°C, or 4.6%), and with higher incoming radiation (by 5.9%), as shown in Supplementary Fig. 3. Nevertheless, intra-seasonal differences also reveal that other factors (e.g., water and sediment conditions, prevalence of growth form) should be taken into account to explain *Ludwigia* ecotype specialisation (Gérard et al., 2014; Thouvenot et al., 2013). Interestingly, other *Ludwigia* photophysiological parameters (α, F_v_/F_m_ and qP) did not differ significantly from those of other species; only its qN was slightly lower than the other macrophytes, highlighting the higher efficiency of *Ludwigia* in dissipating excitation energy through photosynthesis.

On the other hand, absolute photosynthesis and biomass accumulation values are not the only features underlying the success of invasive species. The ability to survive, and even to thrive, in very different environmental conditions could be part of invasive species’ advantage, allowing their spread in areas where they have been introduced. Again, when we consider the variability of *Ludwigia*, we found this species to be distinctive, in that its coefficients of variation for ETR_max_ and I_k_ were at least 62 and 7 times higher than any other species respectively (Table 1). This extreme plasticity could help *Ludwigia* to survive in highly variable conditions encountered in both its native range and in areas to which it is introduced. The other species showing high photophysiological variability was *Trapa* (in terms of F_v_/F_m_, qP, qN), showing its ecological advantage in the Mantua lakes system and explaining its spread in the study area (i.e., nearly half of the total macrophyte-covered surface). It is evident that, in the end, these traits of *Ludwigia* or *Trapa* would largely determine the structure and diversity of the Mantua lakes system macrophyte communities. The results also showed that higher plasticity in a community could be an important determinant of the diversity of a given species. This plasticity helps the species to overcome environmental factors that might otherwise limit its spread in the given environment (Andersen et al., 2012; Lankau, 2011). Not only does the lower mortality at population level undoubtedly result in either higher density and coverage or spatial spreading, but the high variability of functional traits is considered to be an advantage for species establishment and survival (Lankau, 2011; Levine and HilleRisLambers, 2009).

It is also worth comparing the photophysiological responses of *Ludwigia* and *Trapa* to the anomalous meteorological conditions of 2017. Both *Ludwigia* and *Trapa* showed consistent responses to such environmental conditions, in terms of F_v_/F_m_ and qP, with a tendency towards slightly lower responses in 2017 than in 2016. Inversely, the two species reacted differently to the dry and hot weather of 2017, in terms of non-photochemical quenching (qN); compared to July 2016, *Trapa* samples measured in July 2017 tend to have higher qN, while *Ludwigia* had lower levels of non-photochemical quenching in 2017 than in 2016. The reaction of *Ludwigia* to anomalously dry and hot conditions could in part explain the competitive success and invasiveness of water primrose in temperate areas of Europe.

Another key feature facilitating invasive plants in their new environments is the lack of top-down pressure (specific herbivores), and thus the absence of structural defence biomass (Feng et al., 2009; Rout and Callaway, 2009). This results in the preferential allocation of N from structural biomass into leaf tissues and photosystems in invasive plants, consequently increasing the non-structural, photosynthetically active biomass. Our results support this theory, since both *Ludwigia* and *Nelumbo* had significantly higher pigment content (by at least 2-3 times) compared to the native species. The intensification of photosynthetically active biomass helped the plants to further increase the resources available to them, and consequently to extend their presence in their new areas. Besides having the highest absolute values, *Nelumbo* and *Ludwigia* also showed the most variable pigment content, suggesting that these invasive species could adapt to very different environmental conditions. The high pigment content (especially total chlorophyll) of the leaf tissues also affected the specific reflectance of the invasive species and thus the SIs.

PRI was developed and its effectiveness as a proxy for RUE (Gamon et al., 1992; Garbulsky et al., 2011) was demonstrated. Recent works and meta-analysis have shown that PRI also correlates to the seasonal dynamics of pigments pools, and in particular to the ratio of chlorophylls to carotenoids (Filella et al., 2009; Gitelson et al., 2017), and to the daily variability of photosynthetic rates (Filella et al., 2009; Gamon et al., 2015). The PRI literature mainly focusing on terrestrial plants and crops (Garbulsky et al., 2011), not much is known on its ranges and sensitivity for aquatic plants. Our results suggest that, among the macrophyte species considered, the invasive *Nelumbo* might have a higher RUE compared to the native species considered (Fig. 6A). This is possibly due to its higher chl/car ratio (Fig. 4E). PRI_515_, a modification of the original PRI concept and demonstrated to be inversely related to total carotenoid content in terrestrial plants (Hernández-Clemente et al., 2011), shows some notable differences from PRI for allochthonous macrophytes. *Nelumbo* in particular has significantly lower PRI_515_ scores, as well as higher pigment (chlorophylls and total carotenoids; Fig. 4A-C) contents, compared to other species.

Although originally developed for common reed, SIs sensitive to physiological parameters of aquatic plants (Stratoulias et al., 2015) confirm the uniqueness of *Nelumbo* compared to the other species, reinforcing what was already observed for PRI. This species is statistically different from all the others in terms of ND_546,551_, indicating higher chlorophyll fluorescence levels in light-adapted state, and from all the native species in terms of ND_621,692_, which can be attributed to lower values of instantaneous ETR. Even if not statistically significant at the 95% confidence level, *Ludwigia* ND_621,692_ scores are on average slightly lower than those of native species. Additionally, *Nelumbo* mND_705_ scores, related to leaf chlorophyll content in terrestrial plants (Sims and Gamon, 2002), but not yet tested on aquatic vegetation, are significantly higher than any of the other macrophytes. This is similar to the leaf pigment measurements made on our samples (Fig. 4A-C).

The relationship between mND_705_ and pigment contents for macrophyte samples seems to be different for autochthonous and allochthonous species (Supplementary Fig. 4). Significant differences between the species were observed, but *Ludwigia* mND_705_ is not very sensitive to actual pigment contents. From our results, mND_705_ could be effectively used as a proxy for total chlorophyll content for macrophytes, but some caution should be taken (i.e., with the exception of *Ludwigia*, and possibly calibrating a specific model for *Nelumbo*, different from that for *Trapa, Nuphar*, and *Nymphaea*).

Overall, the observed photophysiological and spectral reflectance parameter features point to some notable difference in photosynthetic performance of autochthonous and allochthonous species, either in terms of adaptation to anomalous environmental conditions and maximum photosynthetic rates, in the case of *Ludwigia*, or pigment pool size and higher RUE for *Nelumbo*. In addition, seasonal dynamic features (Table 2) show that both *Nelumbo* and *Ludwigia* have marked differences in season length with respect to *Trapa* and Nymphaeids, with the end of season delayed by approximately 40-60 days (i.e., until November), which arguably constitutes an advantage in terms of productivity.

For *Nelumbo*, this might be related to peculiarities in circadian clock family genes, making it easy for this species to adapt to a wide range of climates and day length regimes, with prolonged flowering times (Ming et al., 2013). In addition, morphological and structural traits (bigger leaves, emerging above water in overlapping layers; average LAI_max_=1.69 m^2^ m^-2^) and fast dynamics during the early vegetative phase (May-June) – with LAI_growth_=0.047 m^2^ m^-2^ dd^-1^ – provide an advantage for *Nelumbo* in the environmental conditions of the Mantua lakes system, and contribute to the success of this species, now established here for approximately a century.

The notion that invasive species tend to show higher scores of functional traits, which emerged in a recent meta-analysis (Van Kleunen et al., 2010), was generally confirmed by our results for the macrophyte species investigated, especially when dealing with traits related to physiology (e.g., leaf pigment contents, ETR_max_, RUE), size (LAI_max_), growth rate (LAI_growth_), and fitness (phenology timing, growing season length). *Ludwigia* and *Nelumbo*, though, have been shown to outperform autochthonous plants in different group of traits, with the former having a greater advantage in terms of physiology-related traits, and the latter in terms of size and growth rate-related traits.

In summary, marked differences of *Ludwigia* and *Nelumbo* compared to native macrophytes, in terms of pigment contents (Chl-a, Chl-b, Car), growing season length, and end of season timing were observed. Parallel to this, compared to all other species, *Ludwigia* had the highest photophysiological parameters (ETR_max_, I_k_). Differences were detected in *Nelumbo* compared to all other species, in terms of spectral reflectance features (i.e., PRI, PRI_515_, ND_546,551_, and mND_705_, connected to higher RUE and pigment pool size), as well as morphological and growth rate features. During the anomalously hot and dry 2017 season, invasive species were found to react differently from native species, as well as between themselves. *Ludwigia* performs differently in terms of α, qN, and ND_546,551_, linked to the variability in photosynthetic efficiency, while *Nelumbo* performs differently in terms of PRI_515_ and mND_705_, mainly related to pigment contents and composition. As these parameters are good indicators of enhanced productivity, such an outcome suggests that an increase in temperature, as for current climate change projections, may further favour the spread of invasive species in temperate areas.

## Conclusions

Solar radiation is an essential resource for plants, and whichever species gain ascendancy over its competitors could be dominant. We found that the success of an invasive macrophyte, in terms of persisting and propagating in its new ecosystem, may be the result of multiple ecological strategies employed. The specificities of the new host areas (such as the lack of top-down pressure, lack of pathogens, etc.) reduce defence costs, thus liberating resources to extend and intensify photosynthesis. This, coupled with their better ability to compete for resources and to tolerate harsh conditions, consequently improves the chances of survival of non-native species. *Ludwigia hexapetala* and *Nelumbo nucifera* have been documented to establish, spread, and alter the Mantua lakes system. Our data show that specific performance-based traits of the invasive macrophytes, photophysiological efficiency, pigment pool size and balance, and leaf spectral reflectance specifically, can describe and explain the success of these species over native ones in the same environment, in terms of both resource competitiveness and tolerance to variability in environmental conditions.

## Acknowledgements

This study was funded by grants from the Hungarian National Research, Development and Innovation Office the NKFIH K-116666 and NKFIH KH-129505, and MTA-CNR bilateral grants and agreement on scientific cooperation between the Hungarian Academy of Sciences and the Consiglio Nazionale delle Ricerche (MacroSense project). Part of the work was carried out in the context of the EU FP7 INFORM project (Grant No. 606865). The authors thank the Parco del Mincio authority and voluntary ecological guards for support given during fieldwork on the Mantua lakes system. The authors are grateful to Stephanie C. J. Palmer for her help with the English of the text.

