## Supplementary Materials for "Aspects of invasiveness of *Ludwigia* and *Nelumbo* in shallow temperate fluvial lakes"

**Supplementary table 1.** Spectral indices derived from macrophyte leaf reflectance measurements used in this study.

| Spectral Index | Acronym | Formula | Leaf parameters | Reference |
|---|---|---|---|---|
| Normalized Difference 750/705 | $ND_{705}$ | $\frac{\rho_{750}-\rho_{705}}{\rho_{750}+\rho_{705}}$ | Chlorophylls (a+b) | Sims and Gamon, 2002 |
| modified Normalized Difference 750/705 | $mND_{705}$ | $\frac{\rho_{750}-\rho_{705}}{\rho_{750}+\rho_{705}-2\rho_{445}}$ | Chlorophylls (a+b) | Sims and Gamon, 2002 |
| Chlorophyll Index | CI | $\frac{\rho_{781}}{\rho_{700}}-1$ | Chlorophylls (a+b) | Gitelson et al., 2003 |
| Anthocyanin reflectance index 2 | $ARI_2$ | $\left(\frac{1}{\rho_{550}}-\frac{1}{\rho_{700}}\right)\rho_{800}$ | Anthocyanins, Carotenoids | Gitelson et al., 2001 |
| Photochemical Reflectance Index | PRI | $\frac{\rho_{531}-\rho_{570}}{\rho_{531}+\rho_{570}}$ | RUE, Car/Chl ratio, xanthophyll EPS | Gamon et al., 1992 |
| Photochemical Reflectance Index 515 | $PRI_{515}$ | $\frac{\rho_{515}-\rho_{531}}{\rho_{515}+\rho_{531}}$ | RUE, xanthophyll EPS (minimize canopy structure dependence) | Hernandez-Clemente et al., 2011 |
| Revised Photochemical Reflectance Index 2 | $PRI_{r2}$ | $\frac{\rho_{531}-\rho_{570}}{2\rho_{550}-\rho_{531}-\rho_{570}}$ | RUE | Wu et al., 2010 |
| PRI combination with Chlorophyll Index | PRI*CI | $\frac{\rho_{531}-\rho_{570}}{\rho_{531}+\rho_{570}}\left(\frac{\rho_{781}}{\rho_{700}}-1\right)$ | RUE, Carotenoids | Garrity et al., 2011 |
| Normalized Difference Spectral Index 546,551 | $ND_{546,551}$ | $\frac{\rho_{546}-\rho_{551}}{\rho_{546}+\rho_{551}}$ | Apparent Chl fluorescence in LAS (Fs) | Stratoulias et al., 2015 |
| Normalized Difference Spectral Index 539,560 | $ND_{539,560}$ | $\frac{\rho_{539}-\rho_{560}}{\rho_{539}+\rho_{560}}$ | Maximum Chl fluorescence in LAS (Fm') | Stratoulias et al., 2015 |
| Normalized Difference Spectral Index 659,687 | $ND_{659,687}$ | $\frac{\rho_{659}-\rho_{687}}{\rho_{659}+\rho_{687}}$ | effective quantum yield of photosystem II (Y(II)) | Stratoulias et al., 2015 |
| Normalized Difference Spectral Index 621,692 | $ND_{621,692}$ | $\frac{\rho_{621}-\rho_{692}}{\rho_{621}+\rho_{692}}$ | Electron Transport Rate (ETR) | Stratoulias et al., 2015 |

**Supplementary table 2.** Summary of *in situ* samples collected for this study.

| Date | Location within Mantua lakes system | | | Species | Samples | | |
|---|---|---|---|---|---|---|---|
| | Lake | Lat (N) | Lon (E) | | Fluorescence | Pigments | Leaf spectra |
| 27/07/2016 | Superior | 45.16139 | 10.72444 | *Nelumbo* | 6 | - | 6 |
| 27/07/2016 | Superior | 45.16069 | 10.73469 | *Nuphar* | 6 | - | 6 |
| 27/07/2016 | Superior | 45.16081 | 10.73567 | *Trapa* | 6 | - | 6 |
| 27/07/2016 | Superior | 45.16111 | 10.73306 | *Trapa* | 6 | - | 6 |
| 28/07/2016 | Superior | 45.16238 | 10.70997 | *Ludwigia* | 6 | - | - |
| 29/07/2016 | Middle | 45.17091 | 10.79712 | *Ludwigia* | 3 | - | 3 |
| 29/07/2016 | Middle | 45.16429 | 10.80396 | *Trapa* | 6 | - | 6 |
| 29/07/2016 | Middle | 45.16853 | 10.79289 | *Trapa* | 6 | - | 6 |
| 29/07/2016 | Inferior | 45.15126 | 10.81209 | *Trapa* | 6 | - | 6 |
| 29/05/2017 | Superior | 45.16244 | 10.71012 | *Ludwigia* | 12 | 9 | - |
| 29/05/2017 | Superior | 45.16306 | 10.78186 | *Nelumbo* | 3 | - | 3 |
| 29/05/2017 | Superior | 45.16290 | 10.77687 | *Nelumbo* | 3 | - | 3 |
| 29/05/2017 | Superior | 45.16036 | 10.76704 | *Nelumbo* | 3 | - | 3 |
| 29/05/2017 | Superior | 45.16247 | 10.70917 | *Nuphar* | 12 | 12 | - |
| 29/05/2017 | Superior | 45.16077 | 10.73465 | *Nuphar* | 4 | 4 | 4 |
| 29/05/2017 | Superior | 45.16889 | 10.78694 | *Nuphar* | 4 | - | 4 |
| 29/05/2017 | Superior | 45.16056 | 10.73583 | *Trapa* | 4 | - | 4 |
| 30/05/2017 | Middle | 45.17052 | 10.79286 | *Nymphaea* | 4 | 4 | 4 |
| 30/05/2017 | Middle | 45.16473 | 10.80586 | *Trapa* | 4 | 4 | 4 |
| 30/05/2017 | Middle | 45.16857 | 10.79258 | *Trapa* | 4 | 4 | 4 |
| 30/05/2017 | Inferior | 45.14895 | 10.81461 | *Trapa* | 4 | 4 | 4 |
| 30/05/2017 | Inferior | 45.15147 | 10.81282 | *Trapa* | 6 | 4 | 6 |
| 27/07/2017 | Superior | 45.16271 | 10.70921 | *Ludwigia* | 9 | 9 | 8 |
| 27/07/2017 | Superior | 45.16330 | 10.78280 | *Nelumbo* | 9 | 9 | 6 |
| 27/07/2017 | Superior | 45.16233 | 10.77392 | *Nelumbo* | 6 | 6 | 6 |
| 27/07/2017 | Superior | 45.16109 | 10.76820 | *Nelumbo* | 6 | 6 | 6 |
| 27/07/2017 | Superior | 45.16271 | 10.70921 | *Nuphar* | 3 | 3 | 3 |
| 27/07/2017 | Superior | 45.16325 | 10.74734 | *Trapa* | 9 | 8 | 9 |
| 28/07/2017 | Middle | 45.17044 | 10.79211 | *Nuphar* | 5 | 6 | 6 |
| 28/07/2017 | Middle | 45.17054 | 10.79266 | *Nymphaea* | 6 | 6 | 6 |
| 28/07/2017 | Middle | 45.16519 | 10.80533 | *Trapa* | 6 | 6 | 6 |
| 28/07/2017 | Middle | 45.16880 | 10.79127 | *Trapa* | 6 | 6 | 6 |
| 28/07/2017 | Inferior | 45.14952 | 10.81424 | *Trapa* | 6 | 6 | 6 |

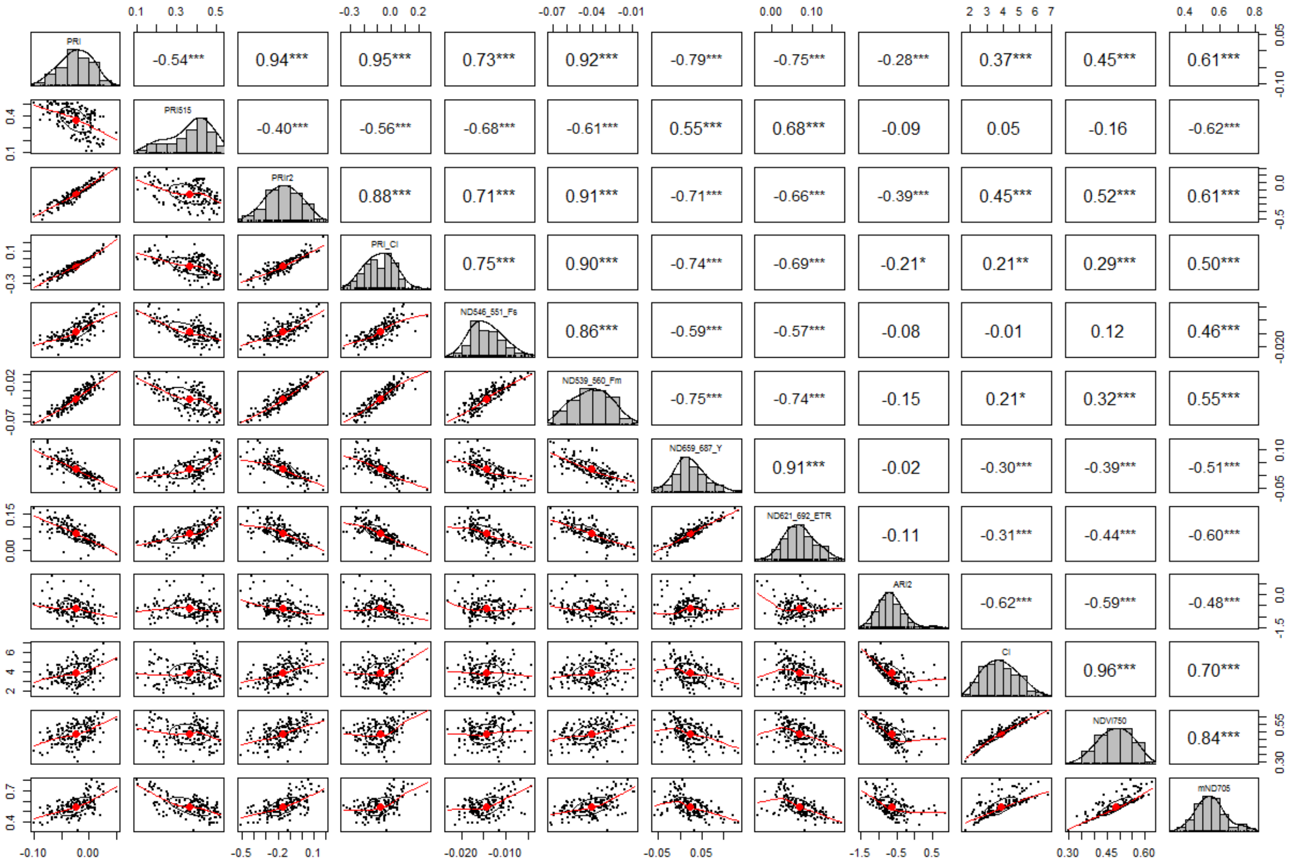


**Supplementary Figure 1.** Correlation matrix of spectral indices (SIs) derived from leaf reflectance measurements of macrophyte samples. Scatter plots of each SI pair are shown in the lower left half; histograms of each SI scores are shown on the diagonal; in the upper right half, the coefficient of correlation for each SI pair (Pearson's r) is shown, together with its significance level (* $p<0.05$, ** $p<0.01$, *** $p<0.001$).

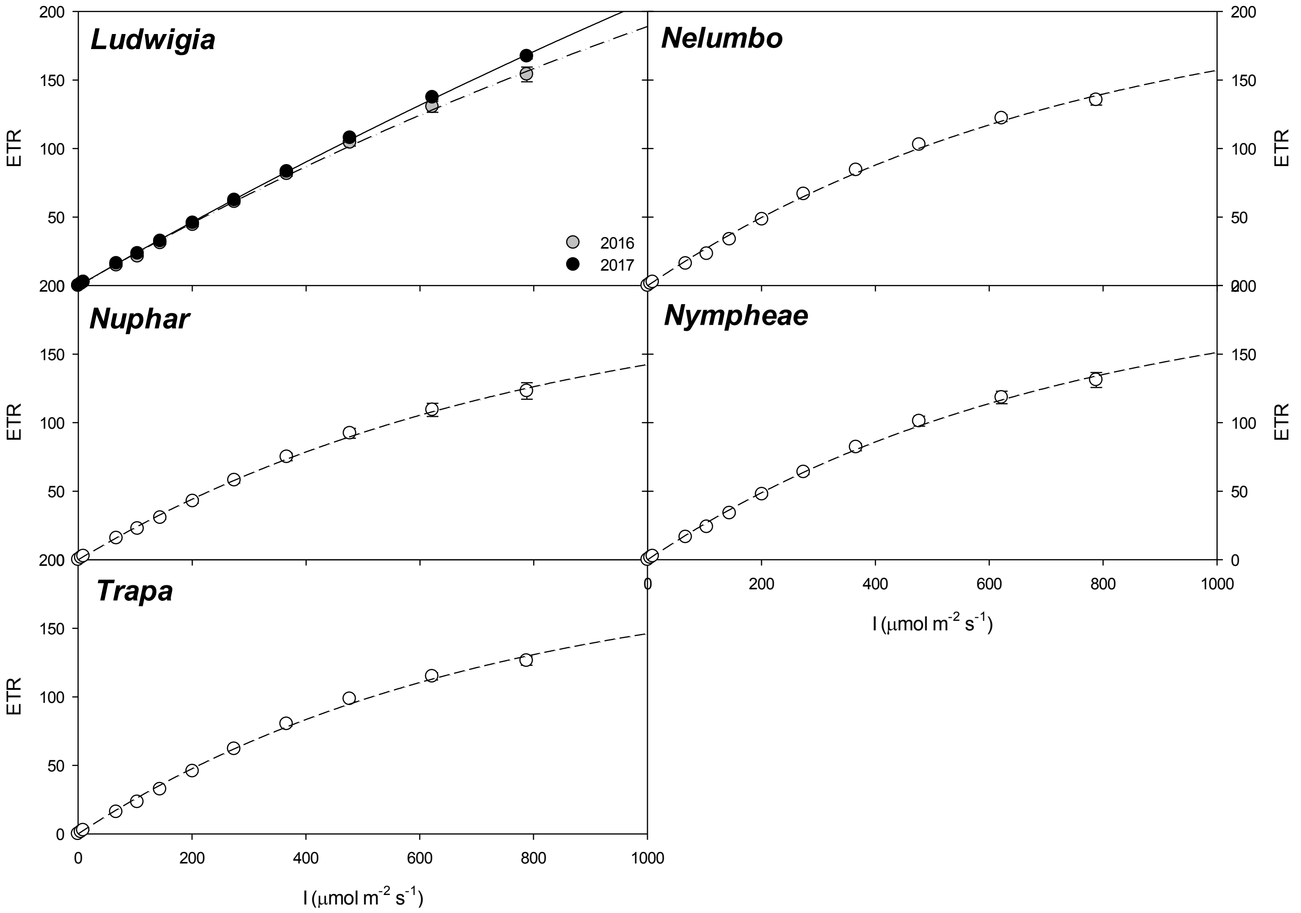


**Supplementary Figure 2.** Effect of light intensity (I) on the electron transport rate (ETR) of the studied macrophytes of the Mantua Lakes system.

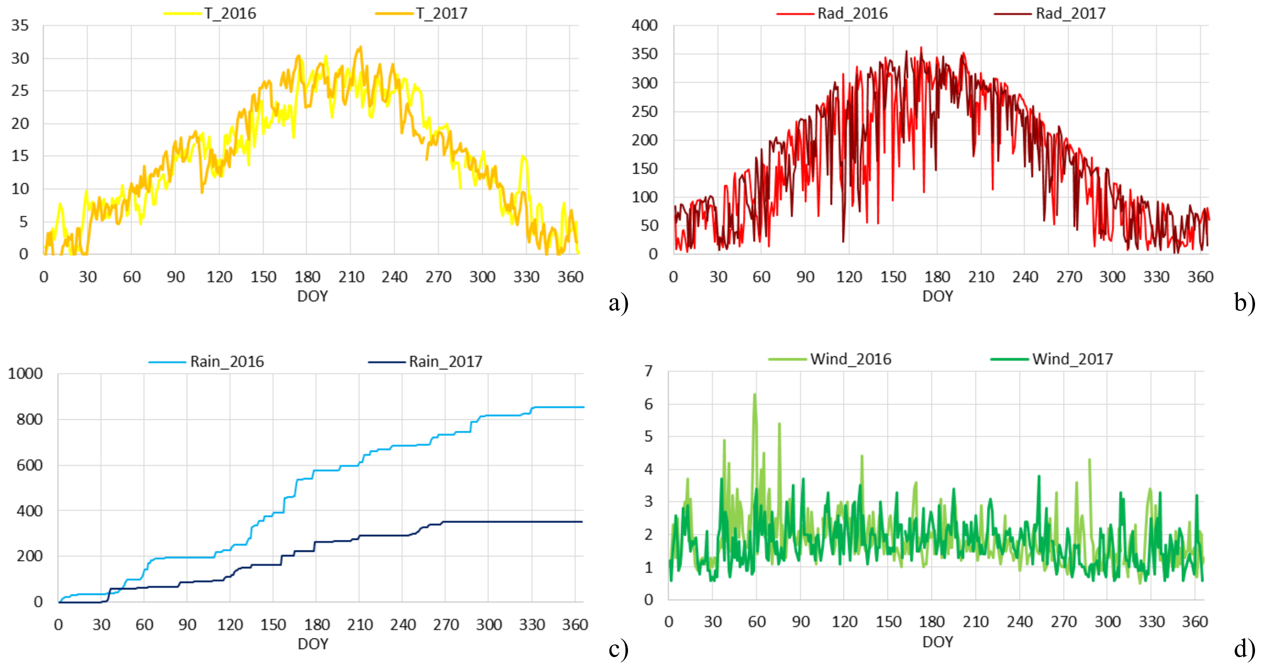


**Supplementary Figure 3.** Time series of meteorological parameters collected during 2016 and 2017 near the study area (45°09’06’’ N; 10°50’04’’ E): a) average daily air temperature (T; °C); b) average daily global incoming radiation (Rad; W $m^{-2}$); c) cumulative precipitation from beginning of the year (Rain; mm); and d) average daily wind velocity (Wind; m $s^{-1}$). Data collected by the Environmental Protection Agency of Lombardy region (ARPA Lombardia), weather station ID: “Mantova Lunetta2 SMR”.

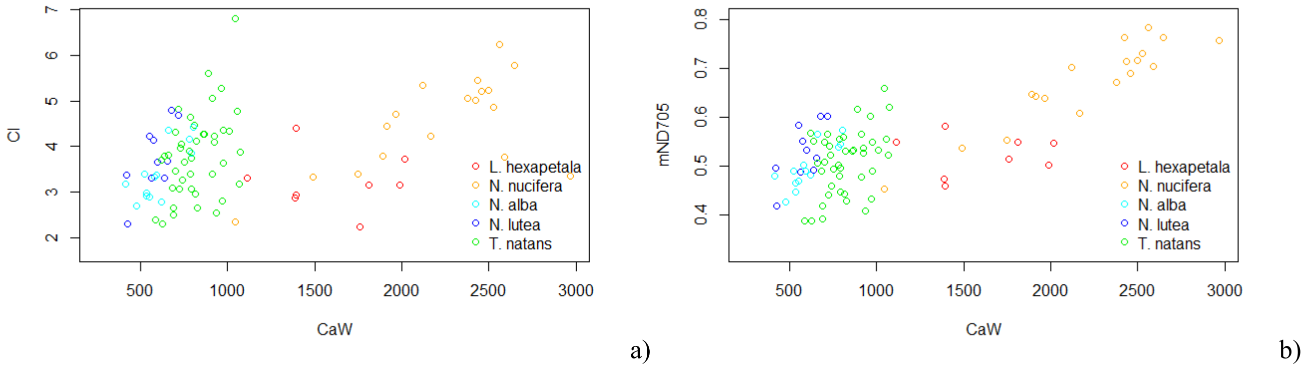


**Supplementary Figure 4.** Leaf Chl-a content (CaW; µg $g_{fw}^{-1}$) versus spectral indices found to be sensitive to chlorophyll content, developed and tested for terrestrial plants, and calculated here for the macrophyte species sampled: a) Chlorophyll Index (CI; Gitelson et al., 2003); b) modified Normalized Difference 750/705 ($mND_{705}$; Sims and Gamon, 2002).

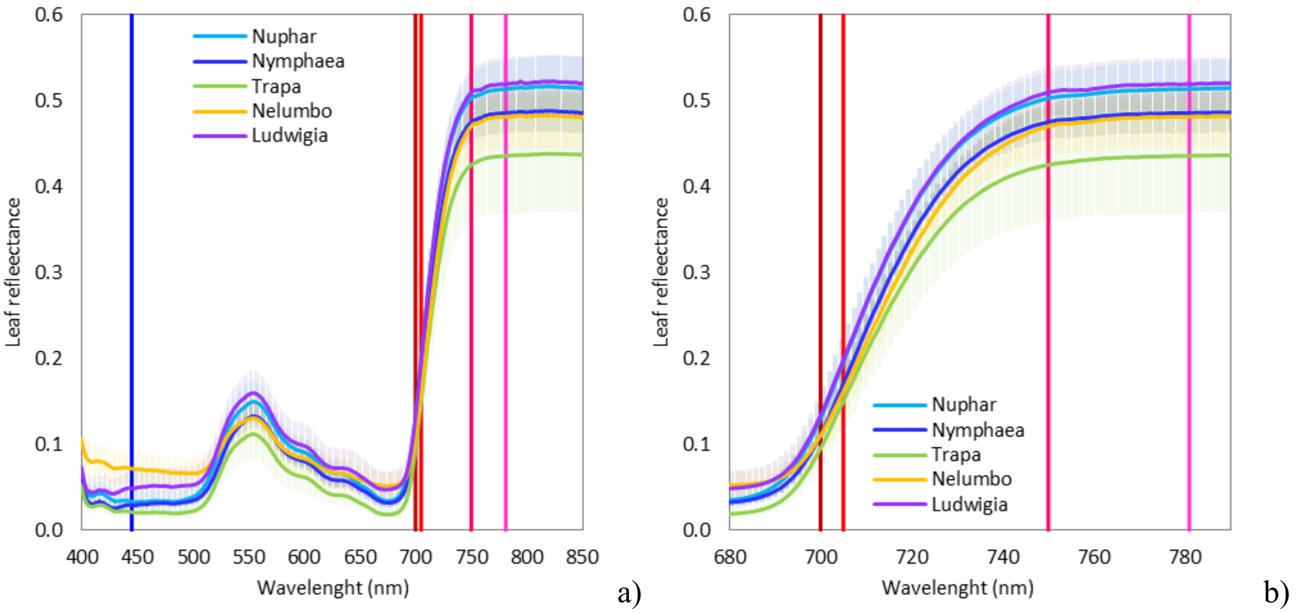


**Supplementary Figure 5.** Leaf spectral reflectance for sampled macrophyte species (average and standard deviation across all samples of each species): a) spectral response in visible to near infrared (VNIR) range; b) detail of spectral response across the red edge range. The plots include marks (vertical lines) at singular wavelengths used in the formulas of Spectral indices sensitive to chlorophyll content ($ND_{705}$, $mND_{705}$, CI). Compared to allochthonous species, the spectral response of *Nelumbo* and *Ludwigia* show specific features of in the red edge to near infrared range (around 690-790 nm), that could be attributed to the differences in mesophyll structure and dry matter: SLA values for these two species are in fact significantly lower than those of native ones. Differences in terms of $mND_{705}$ and chlorophyll content between *Ludwigia* and *Nelumbo,* highlighted in Supplementary Figure 4, could be attributed to peculiar reflectance features in the red edge spectral range (690-730 nm), where the average difference of leaf reflectance for *Ludwigia* and *Nelumbo* is quite stable (around 4-5%) and higher than what should be expected from chlorophyll content alone. The better performance in capturing general variability of leaf chlorophyll content in macrophyte samples by $mND_{705}$ ($r$=0.77) compared to CI ($r$=0.41) reveals the importance of taking into account structural properties and superficial effects (e.g. due to waxes) typical of aquatic plant leaves, which are expressed at shorter visible wavelengths (such as 445 nm included in calculation of $mND_{705}$).
